# A filamentous phage triggers antiviral responses in cystic fibrosis basal airway epithelial cells

**DOI:** 10.1101/2022.09.27.509771

**Authors:** Medeea C Popescu, Naomi L Haddock, Elizabeth B Burgener, Laura S Rojas-Hernandez, Gernot Kaber, Aviv Hargil, Paul L Bollyky, Carlos E Milla

## Abstract

Basal airway epithelial cells are a multipotent stem cell population which gives rise to several airway cell types. Basal cells are known to be critical to airway epithelium homeostasis and repair, and altered basal cell phenotypes have been reported in cystic fibrosis and idiopathic pulmonary fibrosis. However, very little is known about how basal cells respond to stimuli in the cystic fibrosis airway environment. Cystic fibrosis patients experience chronic infection with *Pseudomonas aeruginosa* and contain both high quantities of lipopolysaccharide (LPS) as well as the filamentous bacteriophage Pf produced by biofilm-state *P. aeruginosa* in the airway. In this study, we sought to investigate the transcriptional responses of human basal cells from both healthy controls and patients with cystic fibrosis to LPS and Pf phage. Basal cells from wildtype and cystic fibrosis donors were cultured *in vitro* and exposed to LPS and/or Pf phage, followed by single-cell sequencing on the 10x platform. We report that basal cells show strong antiviral responses and neutrophil chemokine production in response to Pf phage. We validate these findings in additional donors by qRT-PCR and show that Pf phage is internalized by basal cells. We also show that Pf decreases basal cell migration and proliferation. We demonstrate that Pf phage, a bacteria-infecting virus which does not replicate in mammalian cells, is taken up by basal cells and activates immune responses. Further studies are needed to determine the impact of this antiviral response to bacterial clearance.

**Author Summary:** When we experience a lung infection or injury, the stem cells of the airway—called basal epithelial cells—are crucial for healing the affected tissue. Basal cells proliferate and migrate to close gaps in the epithelium and replace injured and dead cells. We are also learning that basal cells can directly contribute to the immune response against lung pathogens, although little is understood about how basal cells sense viruses and bacteria, and what molecules they secrete in response. In our work, we sought to investigate the role that basal cells play in the course of an infection with the common human pathogen *Pseudomonas aeruginosa*. We stimulated basal cells with lipopolysaccharide (LPS), a potent immunogen produced by *Pseudomonas*, as well as the *Pseudomonas-*infecting virus Pf4. We have previously shown that Pf4, which is produced in high amounts by lung-infecting Pseudomonas, affects the immune response to this bacterium. In our work here, we show that Pf4 is recognized as a virus by basal cells and changes the transcriptional response to LPS, derailing the antibacterial immune response to *Pseudomonas*. These findings raise the question of whether other bacteria-infecting viruses alter immune responses, and how these viruses interact with other airway cells.

## Introduction

Basal respiratory epithelial cells are a stem cell population playing critical roles in homeostasis and repair of airway epithelium. These cells make up approximately 30% of the pseudostratified mucocilliary epithelium of the tracheobronchial tree and are present throughout the airways, differentiating to regenerate key airway cell types[1]. They are critical in injury response and in repair of airway tissue[2]. These epithelial cells provide a barrier to pathogens in the lung and contribute to innate responses. Basal cells possess Toll-like receptors for recognition of pathogen-derived molecules and are capable of secreting inflammatory mediators, chemokines, and antimicrobial substances in response to bacterial stimulus[3,4]. Previous reports in explanted lung tissue from end-stage lung disease in cystic fibrosis (CF) and idiopathic pulmonary fibrosis indicate that basal cells are a heterogenous cell type, with altered phenotypes [5,6]. Basal cells are of interest for their roles in chronic respiratory disease and contribution to inflammation in the airways of patients with CF.

CF is an autosomal recessive genetic disease caused by mutations in the cystic fibrosis transmembrane conductance regulator (CFTR) gene, leading to mucus buildup and chronic bacterial infections. Most patients with CF are colonized by *P. aeruginosa* (Pa) by adulthood[7]. Pa infection becomes a major contributor to patient outcomes later in life – chronic Pa infection is associated with declining lung function, increased episodes of pulmonary exacerbations, and increased mortality[8–10]. In addition to bacterial burden, our group has shown that patients with CF infected with Pa also harbor high titers of Pa-infecting bacteriophages in the Pf family[11]. We found approximately 40% of patients with CF infected with Pa to also have Pf present in their sputum, with 10^6^-10^11^ Pf virions/ml detected. The impact of this high burden of Pf phage in infected airways is not fully understood.

Pf phages are a genus of filamentous bacteriophages which chronically infect Pa. The phage genome integrates into the Pa genome as a lysogen, and virons are continuously extruded from the bacterial cell envelope without lysis[12–15]. Presence of Pf phages in a Pa biofilm has been shown to contribute to infection persistence and chronicity, and we have also demonstrated that Pf phages engage anti-viral pattern recognition receptors in myeloid cells[16–18]. In patients with CF, Pf carriage is associated with chronic infection, older age, larger decrease in lung function during pulmonary exacerbation, and antibiotic resistance[11]. Our goal is to identify cellular mechanisms for these associations, building on previous work establishing Pf as a virulence factor produced by Pa.

In this work, we asked whether the abundant phages produced in the course of chronic Pa infection affect basal airway epithelial cells by investigating transcriptional responses to Pf phage in the context of bacterial stimulation.

## Methods

### Basal Cell Culture

Nasal brushings were acquired under a human subjects-approved protocol (SU IRB # 42710) from a CF patient with a 1288insTA/406-1G>A genotype, a CF patient with a F508del/F508del genotype and from a healthy control known to be wildtype at the CFTR locus (WT). Samples were procured and processed as described previously [19,20]. Once confluent, cells were treated for 24 h with 5 μg/mL LPS, PBS control, and 1×10^10^ pfu/well of purified phage preparation (an effective MOI of 1000 phage copies per eukaryotic cell). In addition to evaluating responses to Pf4 phage, we included as controls the *E. coli* filamentous phage Fd and the Pa lytic phage DMS3vir. These phage were prepared as described previously [18,21].

### Single-cell Sequencing

Single-cell suspensions of basal cell culture were prepared by lifting adherent cells using trypsin digestion. Dead cells were removed using the MACS Dead Cell Removal kit (Miltenyi Biotech). Cells were then washed and counted as per manufacturer’s instructions and libraries were prepared using the 10X NEXT GEM 3’ Gene Expression Library scRNA Seq v3.1 reagents and workflow (10x Genomics). Library concentration and quality was confirmed by BioAnalyzer.

Libraries were sequenced on the HiSeq PE150 platform. Quality control of raw data was performed using FASTQC, and reads were aligned to the GRCh38-2020 Human Reference Genome using CellRanger to generate counts. An average of 8328 cells were sequenced across all conditions with an average depth of 29,962 reads/cell. Demultiplexed data was analyzed in R using the Seurat 4.0 package[22].

### qRT-PCR

Basal cells from additional WT and CF (F508del/F508del genotype) donors were cultured and treated as described above, and total RNA was extracted using the Rneasy RNA Extraction kit (QIAGEN). RNA was used to synthesize cDNA through reverse transcription (Applied Biosystems). qRT-PCR was performed to quantify antiviral gene expression. Primers sequences were obtained from Primerbank and previously published work, and were as follows: GAPDH[23], OAS1 (Primerbank #74229012c1), OAS2 (Primerbank #74229020c1), IRF7 (Primerbank #98985817c1), and IFIT3 (Primerbank #197276657c1). Three independent validation experiments were performed for CF cells, and two independent validations were done for WT cells.

### Microscopy

Basal cells were cultured on collagen-coated cover slips for 48 h and then treated with 10^11^ pfu/mL purified Alexa488-labelled Pf phage or PBS control for 24 h. Cells were then fixed in 4% paraformaldehyde, permeabilized in 0.1% Triton X-100, stained with primary anti-EEA-1 antibody for 2 h and secondary AF647 antibody for 30 minutes. Cells were then stained with phalloidin-CF555 for 20 minutes, followed by DAPI for 10 minutes. Slides were imaged on a Leica confocal microscope using Leica LASX software at 100x magnification, and resulting images were processed in ImageJ.

### ELISA

Supernatants from basal cell cultures were assessed for cytokine content as per manufacturer’s instructions. CXCL1 (R&D systems), IL-8/CXCL8 kits (R&D systems), CXCL5 (Biolegend) and CXCL10 (Biolegend) kits were used.

### Proliferation/Migration Assay

Basal cells were isolated as described above from a patient with CF with a 1288insA / 406-1G->A genotype and WT donor. Cells were plated onto collagen coated 96-well plates and once actively proliferating were treated with 1 μg/mL LPS along with 10^10^ pfu/mL Pf phage or PBS control. The plate was then placed on an Incucyte incubator (Sartorius Lab Instruments, Goettingen) at 37^·^C and 5% CO2 for real time phase contrast-imaging over a 48 h time span. Conditions were run in quadruplicate, and progression to confluence was assessed by image analysis following the manufacturer instructions.

## Results

### WT and CF basal cells cluster into functionally distinct subsets

Cells from a WT donor and patient with CF harboring 1288insTA/406-1G>A mutations were subjected to unsupervised clustering and visualized by UMAP (Fig 1A). Seven biologically meaningful clusters were identified across donors (Fig 1B). Cells showed a clear response to LPS stimulation: gene set enrichment analysis on PBS vs. LPS-treated cells showed an enrichment of inflammatory response genes in LPS-stimulated conditions in both WT and CF cells (Fig S1A). Interestingly, the most significant gene sets enriched in CF LPS-treated cells as compared to WT LPS-treated cells were all related to unfolded protein responses, indicating that the largest difference in these cells remains the presence of misfolded CFTR, and that CF and WT cells have a common LPS response (Fig S1B).

**Fig 1:**
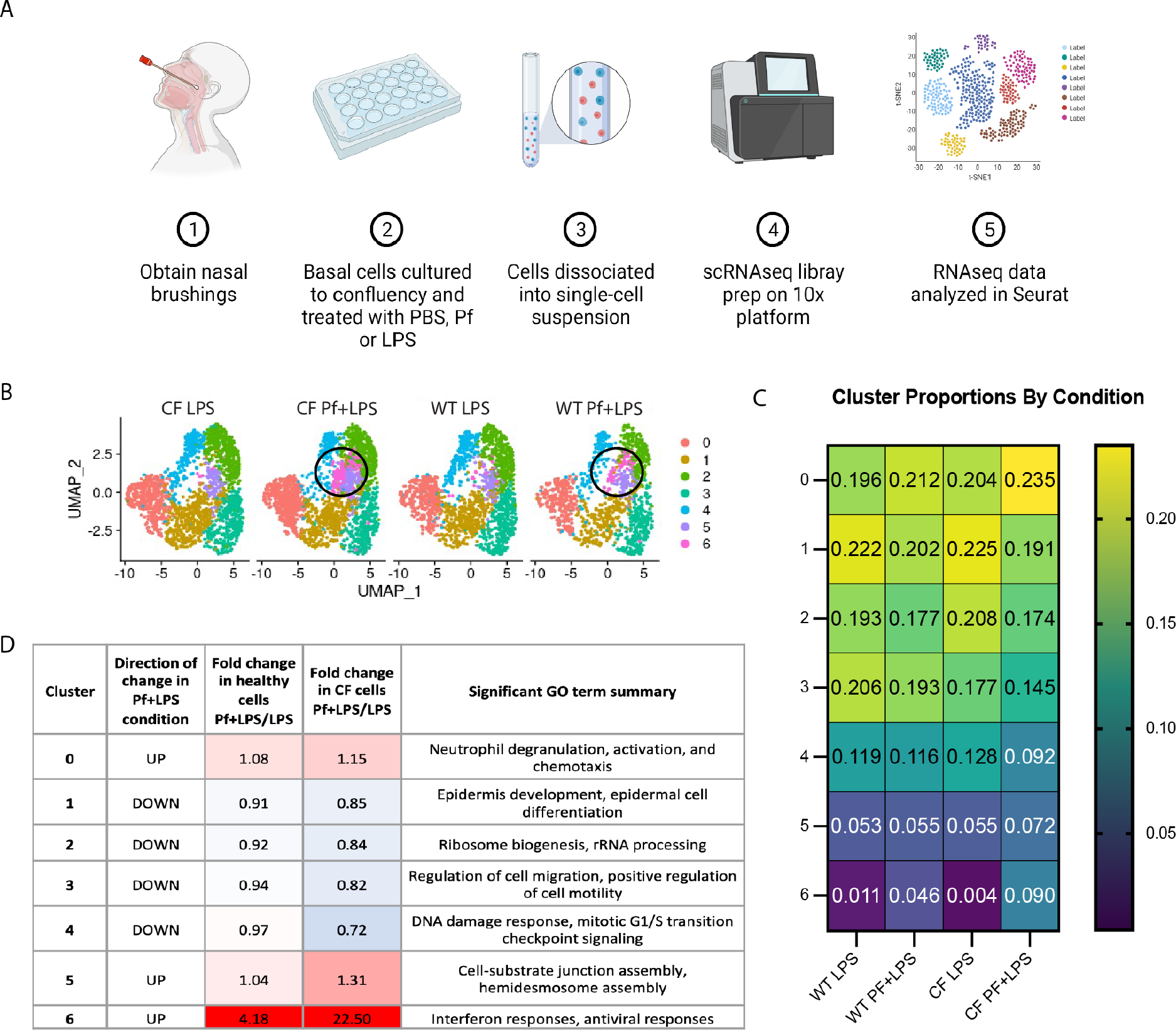
WT and CF primary basal epithelial cells exhibit hetereogenity in response to Pf4 bacteriophage and LPS. (A) Diagram of experimental workflow. (B) UMAP of single-cell sequenced WT basal epithelial cells and CF epithelial cells stimulated with 5 μg/mL LPS ± 10^10^ pfu/mL Pf4 phage for 24 h. Cluster corresponding to antiviral responses circled in black. (C) Proportion of cells in each cluster for WT cells and CF cells between LPS and Pf+LPS conditions. (D) GO terms corresponding to each cluster across genotypes. Graphical schematics created with BioRender.com.

Both CF and WT cells contained a small but significant cluster defined by strong upregulation of antiviral response genes and interferons, which was virtually absent in cells treated with LPS only (Fig 1B-D, Fig S2). This antiviral cluster represented 4.6% of all the cells in the WT donor Pf+LPS condition, and was significantly larger in the CF Pf+LPS condition at 9.0% of cells. The proportion of cells corresponding to this cluster increased ∼4-fold in WT cells, and ∼20-fold in CF cells (Fig 1B-C). A corresponding antiviral cluster was enriched in controls treated with Pf only (Fig S3A-C). Apoptotic markers were not increased in the Pf+LPS conditions and did not correlate with the antiviral cluster marker genes IFI6 or ISG15 (Fig S4A-B), and feature plots of immune cell markers show little to no cells contaminating these samples (Fig S5). Together, these data indicate that this antiviral signature is due to a direct response of these cells and not driven by contaminating or apoptotic cells.

Apart from cluster 6, several other clusters showed changes in cell proportion between conditions. Specifically, following Pf+LPS treatment, both genotypes had lower proportions of cells in clusters related to epidermal differentiation, regulation of cell migration, and DNA damage response/mitotic cell cycle transitioning (Fig 1B-D, Fig S2). These findings indicate that the presence of Pf appears to hinder or divert basal cells from homeostatic and infection/injury-repair processes. Notably, CF cells had larger decreases in these clusters than WT cells. In addition, both WT and CF also showed an increase in the cluster defined by neutrophil activation/chemotaxis, suggesting that the presence of Pf increases the number of cells expressing chemotactic factors as compared to LPS stimulation alone.

### Pf phage induces an antiviral response in a subset of basal cells

In order to further characterize the Pf-driven antiviral cluster, we identified the top-expressed individual genes unique to this subgroup of cells. These sets of genes were virtually identical between WT and CF, and included such canonical type I interferon-driven genes as DDX-, OAS-, and ISG-family genes (Table 1, Fig 2B). Top GO terms for this gene set included cellular response to type I interferon, negative regulation of viral genome replication, and cellular response to dsRNA (Fig 2A). This last enrichment was particularly interesting given our previous findings that Pf phage triggers TLR3, a dsRNA sensor [18]. Furthermore, although the antiviral cluster exhibited the highest expression of antiviral genes, several of these genes were also expressed to varying degrees across all clusters. Notably, overall expression of the top-enriched antiviral genes was higher in the Pf+LPS condition as compared to the LPS-only condition for both genotypes (Fig 2C-D). These antiviral genes were shown to be upregulated in Pf only controls as well, indicating the signature is independent of LPS stimulus (Fig S3D).

**Table 1:** Top enriched genes in cluster 6.

| Gene | P-value | Avg.<br>log(fold<br>change) | Adj. p-value |
| --- | --- | --- | --- |
| ISG15 | 0 | 4.04 | 0 |
| CCL5 | 0 | 1.61 | 0 |
| OASL | 0 | 3.59 | 0 |
| DDX58 | 0 | 2.20 | 0 |
| IFIT1 | 0 | 3.12 | 0 |
| RSAD2 | 0 | 2.47 | 0 |
| IFIT2 | 0 | 3.38 | 0 |
| HERC5 | 0 | 2.04 | 0 |
| IFIH1 | 0 | 1.88 | 0 |
| DDX60 | 0 | 2.36 | 0 |
| IFIT3 | 0 | 3.18 | 0 |
| IFI44 | 0 | 1.83 | 0 |
| ZC3HAV1 | 0 | 2.24 | 0 |
| USP18 | 0 | 1.21 | 0 |
| ISG20 | 0 | 2.03 | 0 |
| CXCL11 | 0 | 2.46 | 0 |
| SAMD9 | 0 | 2.20 | 0 |
| OAS3 | 0 | 1.19 | 0 |
| MX1 | 0 | 2.17 | 0 |
| OAS1 | 0 | 1.58 | 0 |
| PMAIP1 | 0 | 2.55 | 0 |
| OAS2 | 0 | 1.01 | 0 |
| NUPR1 | 0 | 2.42 | 0 |
| PLAUR | 0 | 1.94 | 0 |
| IFI6 | 0 | 1.70 | 0 |
| IFITM3 | 0 | 1.22 | 0 |
| PARP14 | 0 | 1.46 | 0 |
| IRF7 | 0 | 1.11 | 0 |
| EIF2AK2 | 0 | 1.17 | 0 |
| GBP1 | 0 | 1.01 | 0 |

**Fig 2:**
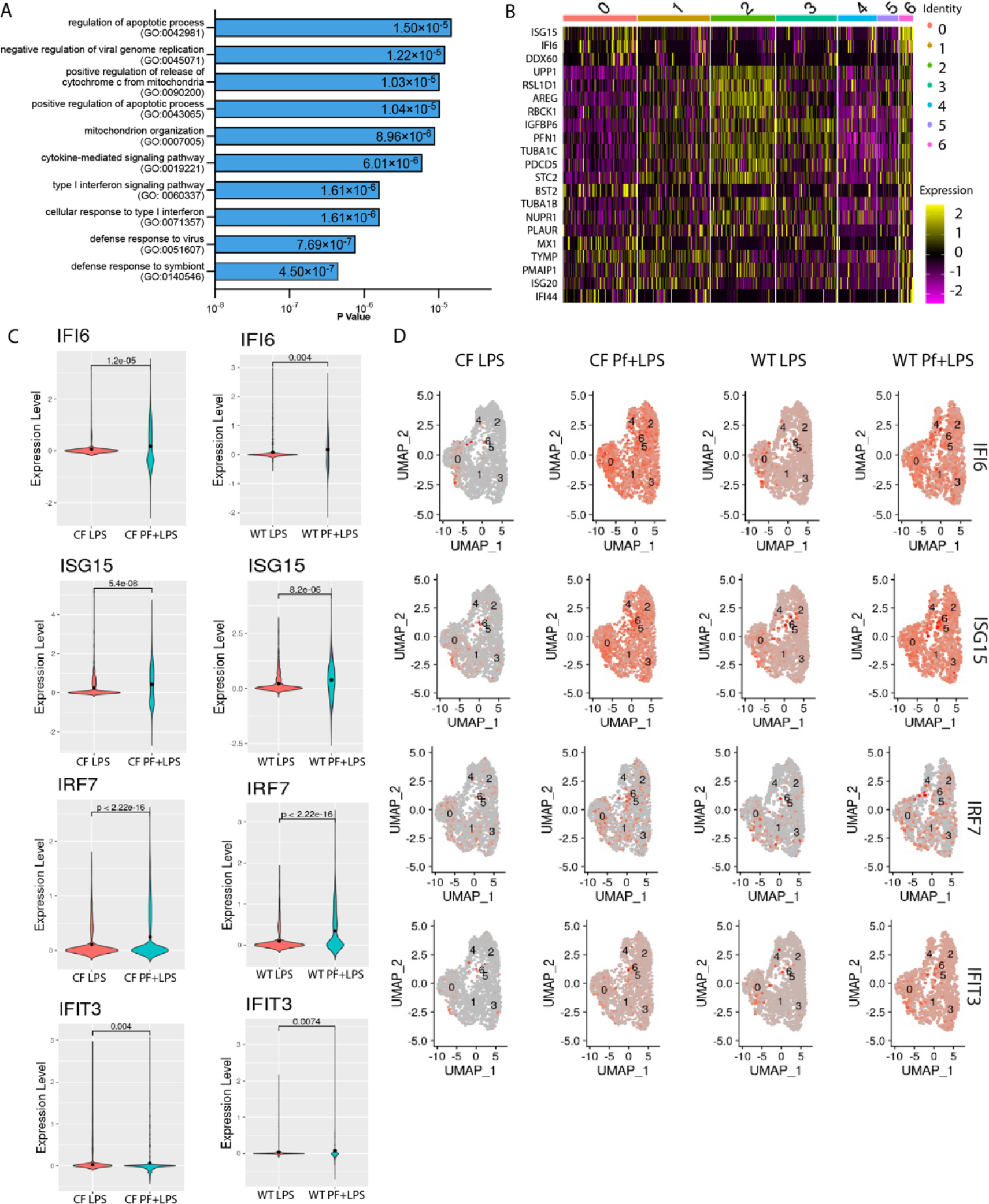
Basal cells initiate antiviral responses to Pf4 phage. (A) GO term enrichment for top upregulated genes in antiviral cluster. (B) Gene expression pattern across all clusters for top upregulated genes of antiviral cluster. (C) Gene expression for individual antiviral genes across control and Pf-stimulated conditions cystic fibrotic (left) and WT (right) cells. (D) Overall expression of individual antiviral genes in WT and CF cells exposed to LPS or Pf+LPS.

We validated the upregulation of these type I interferon response genes in basal cells from an additional WT donor and additional patient with CF with F508del/F508del genotype by qRT-PCR (Fig 3). We chose validation targets with a range of expression intensities, both top-expressed and lower-expression genes (Fig 3C). We found that, as expected, exposure to Pf phage both with and without additional LPS stimulation induced upregulation of the antiviral genes IFIT3, OAS1, OAS2 and IRF7 at levels equal to or above LPS-only gene induction (Fig 3A-B). In particular, IRF7 was exclusively induced by Pf phage stimulation. CF cells showed a greater induction of antiviral genes than WT cells in response to Pf, with IRF7 in particular being much more highly expressed.

**Fig 3:**
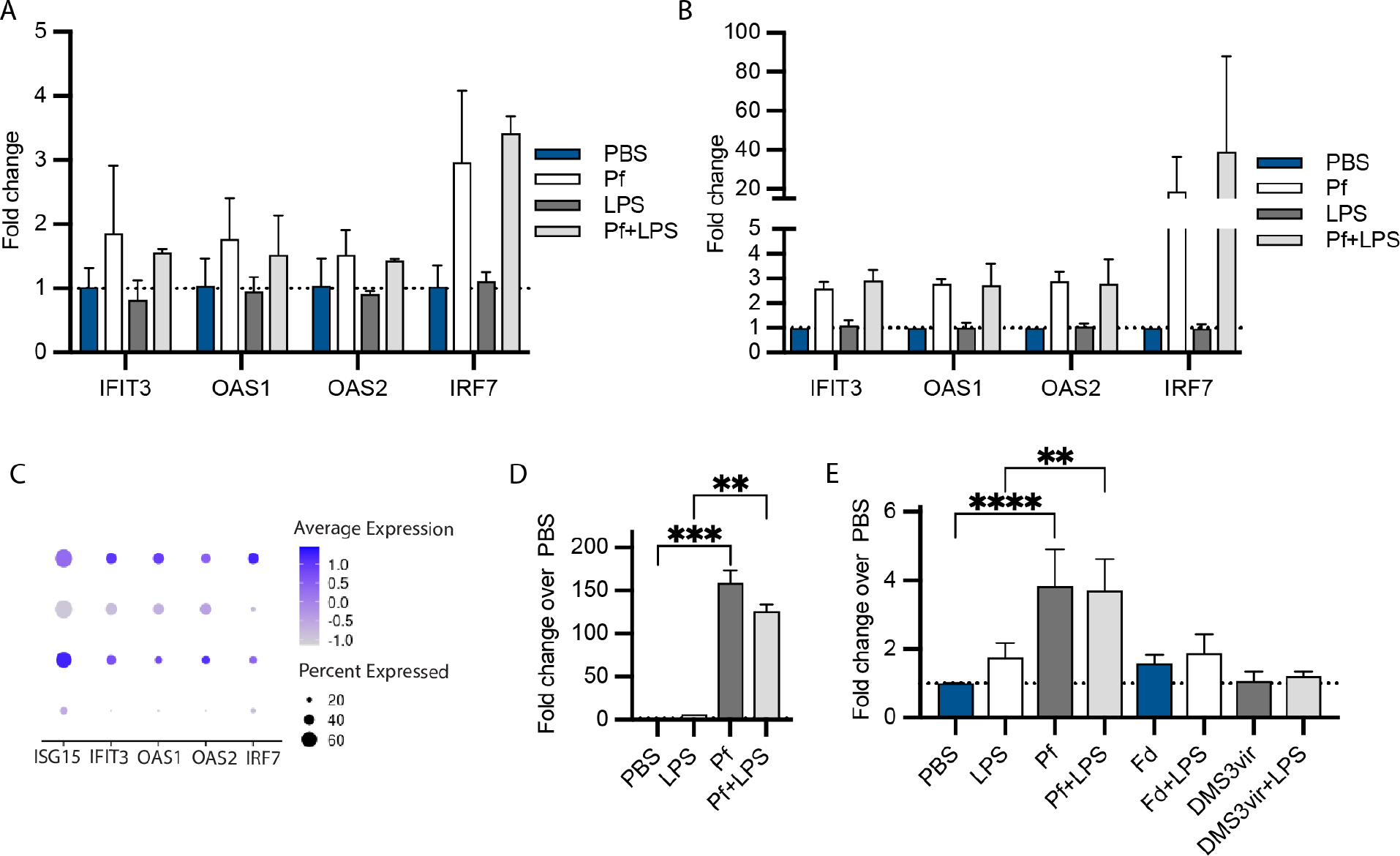
Basal cells from additional donors validate antiviral signature in response to Pf phage. RT-qPCR of WT (A) and CF (B) basal cells incubated with 5 ug/ml LPS ± 10^10^ pfu/mL Pf4 phage for 24hrs. (C) Expression of the selected validation genes in sequencing dataset. Protein quantification of CXCL10 in supernatants from WT (D) and CF (E) basal cells stimulated as in (A) as determined by ELISA, with 10^10^ pfu/mL of additional phage controls. Significance was assessed by ANOVA followed by Holm-Šídák test for multiple comparisons.

We also evaluated basal cell production of the canonical antiviral response chemokine CXCL10. We found that both WT and CF cells produce much higher amounts of CXCL10 in response to Pf stimulation as compared to LPS (Fig 3D-E). Notably, our control phages Fd (produced in *E. coli*) and DMS3vir (produced in Pa) did not induce significant CXCL10 secretion in basal cells (Fig 3E).

We confirmed that induction of Pf-driven antiviral responses could not be driven by LPS or free nucleic acid contamination in our Pf phage preparations. The LPS content of our Pf phage reagent was ∼25 ng/mL at the dilution used, much lower than the 5 ug/mL of LPS used as a stimulus. Effective LPS content of Fd and DMS3vir phages at the dilutions used were 2 μg/mL and 20 ng/mL, respectively, and these phages did not induce antiviral chemokine production. Extensive nuclease digestion is also performed on our phage preparations in order to eliminate free DNA or RNA.

Finally, given that our previous findings implicated a role for TLR3—an endosomal sensor—in sensing Pf phage, we investigated whether basal cells were capable of internalizing phage. Basal cells were cultured *in vitro* as for single-cell sequencing and incubated with 10^11^ pfu/mL Pf phage or PBS control for 24 h. Punctate Pf phage uptake was clearly observed, often overlapping with EEA-1 staining, an early endosome marker, indicating internalization of phage particles (Fig 4).

**Fig 4:**
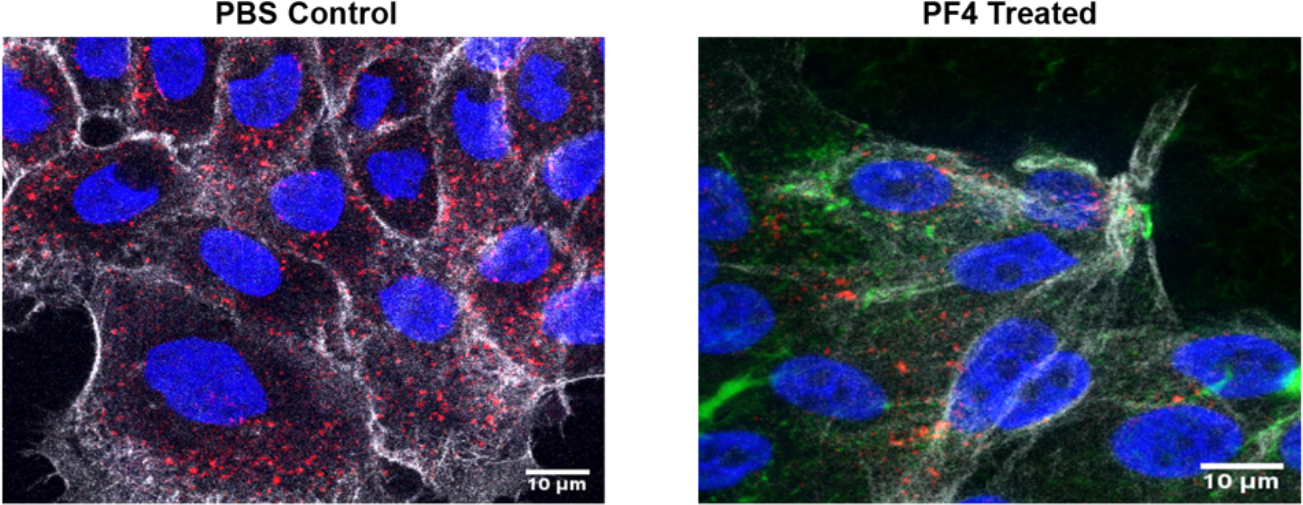
CF basal cells internalize Pf4 phage. Basal cells from one CF donor were expanded *ex vivo* and imaged at 100x magnification. Cells were incubated with 10^11^ pfu/mL of AlexaFluor-488-labelled Pf4 phage for 24 hours, and stained with phalloidin-CF555, EEA-1-AF647, and DAPI.

### Pf phage increases neutrophil chemoattractant production in basal cells

We also chose to further investigate Pf-mediated effects on basal cell chemokine expression. Cluster 0, defined by neutrophil activation and chemotaxis gene upregulation, increased by 10-15% with Pf and LPS stimulation as opposed to LPS-only treated cells in both WT and CF basal cell cultures (Fig 5A, Table 2). This subset of cells expressed genes such as CXCL1, CXCL2, and IL-8/CXCL8, all potent neutrophil migration and activation factors (Fig 5B). We evaluated protein production of a subset of these factors in basal cell cultures from additional WT and CF donors as described above, and confirmed that Pf stimulation strongly increases production of CXCL1, CXCL8, and CXCL5 above the level induced by LPS alone in both genotypes (Fig 5B-D). These markers were also found to be upregulated in Pf only conditions, indicating this effect is independent of LPS stimulus (Fig S3E).

**Table 2:** Top enriched genes in cluster 0.

| Gene | P-value | Avg. log(fold change) | Adj. p-value |
| --- | --- | --- | --- |
| WFDC2 | 0 | 3.55 | 0 |
| SCGB1A1 | 0 | 3.31 | 0 |
| PSCA | 0 | 3.23 | 0 |
| VMO1 | 0 | 3.04 | 0 |
| SAA1 | 0 | 2.98 | 0 |
| PIGR | 0 | 2.94 | 0 |
| MSMB | 0 | 2.85 | 0 |
| C3 | 0 | 2.83 | 0 |
| SERPINA3 | 0 | 2.78 | 0 |
| MUC1 | 0 | 2.72 | 0 |
| LCN2 | 0 | 2.71 | 0 |
| CP | 0 | 2.70 | 0 |
| SLPI | 0 | 2.56 | 0 |
| CAPN13 | 0 | 2.50 | 0 |
| IGFBP3 | 0 | 2.35 | 0 |
| SAA2 | 0 | 2.31 | 0 |
| MUC4 | 0 | 2.30 | 0 |
| SLC6A14 | 0 | 2.17 | 0 |
| RARRES1 | 0 | 2.16 | 0 |
| RDH10 | 0 | 2.14 | 0 |
| CXCL17 | 0 | 2.12 | 0 |
| LYPD2 | 0 | 2.07 | 0 |
| CFB | 0 | 2.03 | 0 |
| GSN | 0 | 2.02 | 0 |
| S100P | 0 | 2.01 | 0 |
| STEAP4 | 0 | 1.96 | 0 |
| C15orf48 | 0 | 1.90 | 0 |
| CXCL1 | 0 | 1.88 | 0 |
| MUC20 | 0 | 1.86 | 0 |
| SERPINB3 | 0 | 1.84 | 0 |
| MUC16 | 0 | 1.84 | 0 |
| IFI27 | 0 | 1.81 | 0 |
| ATP1B1 | 0 | 1.79 | 0 |
| ST6GALNAC1 | 0 | 1.69 | 0 |
| MDK | 0 | 1.68 | 0 |
| TNFSF10 | 0 | 1.67 | 0 |
| CEACAM5 | 0 | 1.66 | 0 |
| GDF15 | 0 | 1.64 | 0 |
| AGR2 | 0 | 1.64 | 0 |
| CEACAM6 | 0 | 1.60 | 0 |
| GOLM1 | 0 | 1.58 | 0 |
| TNFAIP2 | 0 | 1.56 | 0 |
| S100A4 | 0 | 1.52 | 0 |
| CXCL8 | 0 | 1.48 | 0 |

**Fig 5:**
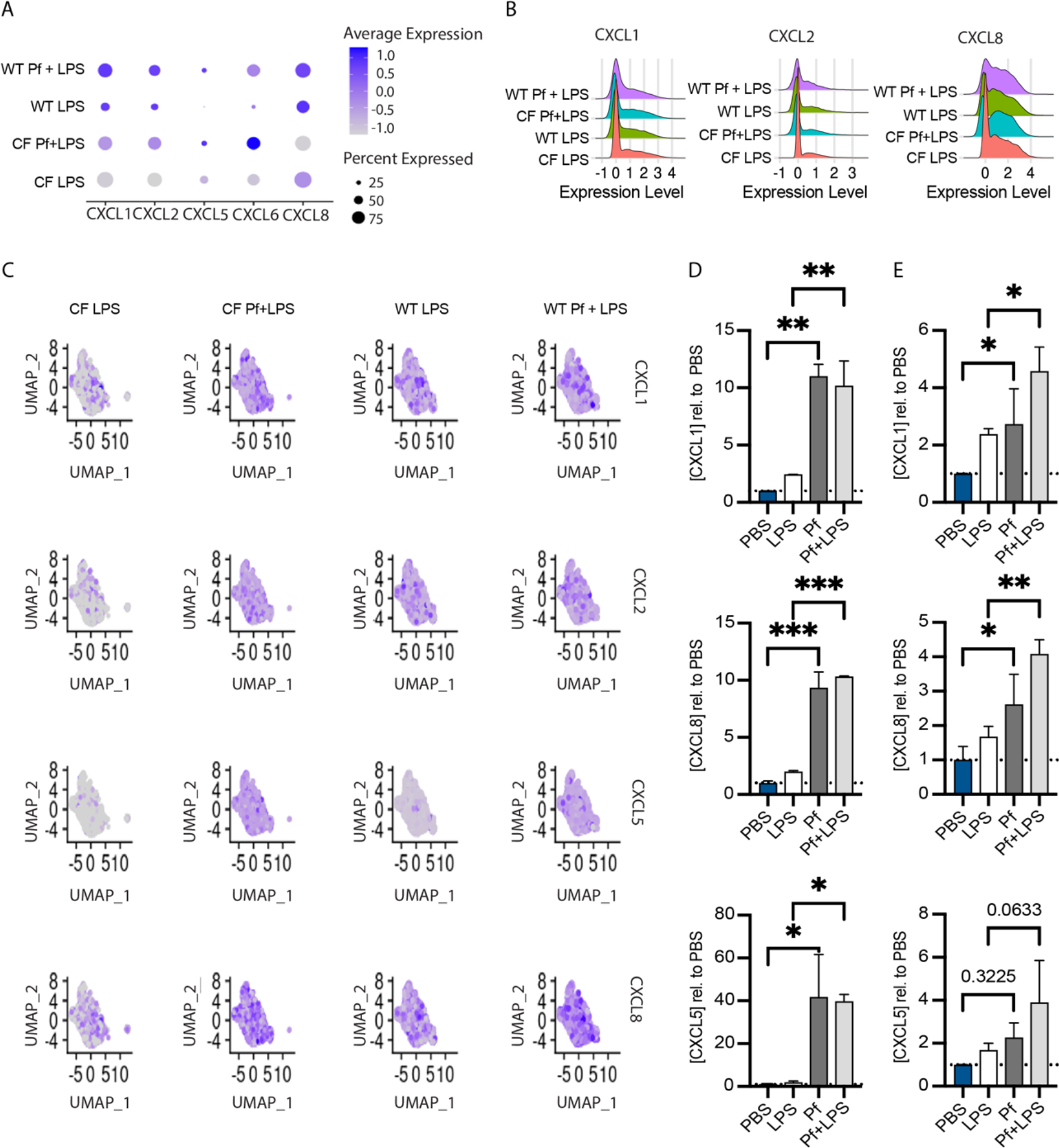
Pf4 phage induces production of neutrophil chemotactic factors in WT and CF basal cells. Expression (A) and ridge plots (B) of selected neutrophil chemoattractant genes in 10x sequencing dataset. (C) Feature plots showing neutrophil chemokine expression across basal cell clusters. Protein concentration of neutrophil chemoattractants in supernatants from WT (D) and CF (E) basal cells incubated with 5 ug/ml LPS ± 10^10^ pfu/mL Pf4 phage for 24 h as determined by ELISA. Significance was assessed by ANOVA followed by Holm-Šídák test for multiple comparisons.

### Pf impairs the ability of basal cells to reach confluence

A final Pf-induced phenotype we chose to investigate was the effect of this phage on the migration and proliferation of basal cells. We noted that a subset of basal cells in both the WT and CF datasets (cluster 3) expressed cellular migration and motility-related genes. This cluster consisted of ∼20% of all basal cells, and was decreased by 6% and 18%, respectively, in WT and CF cells exposed to Pf phage (Fig 1B-C). Although there were no significant differences in cell cycle state between conditions in cells of either genotype in our single-cell dataset, this dataset consisted of confluent cells which had moved past active proliferation and migration. Therefore, in order to further understand the effects of Pf phage on basal cell growth towards confluence, we tracked cellular growth in sparsely seeded, actively proliferating basal cells over the course of 48 h.

Phage treatment appeared to delay progression towards confluence, with a stronger phenotype in CF cells as compared to WT cells (Fig 6A-D). After 48 h untreated CF cells had significantly lower confluence that WT cells (Fig 6E). WT cells exposed to Pf and LPS had a substantial decrease compared to control conditions, with no significant difference between Pf and LPS treatment alone but with their combination inducing the most pronounced significant effect (Fig 6E). CF cells treated with Pf and LPS had the lowest confluence compared to control conditions (Fig 6E).

**Fig 6:**
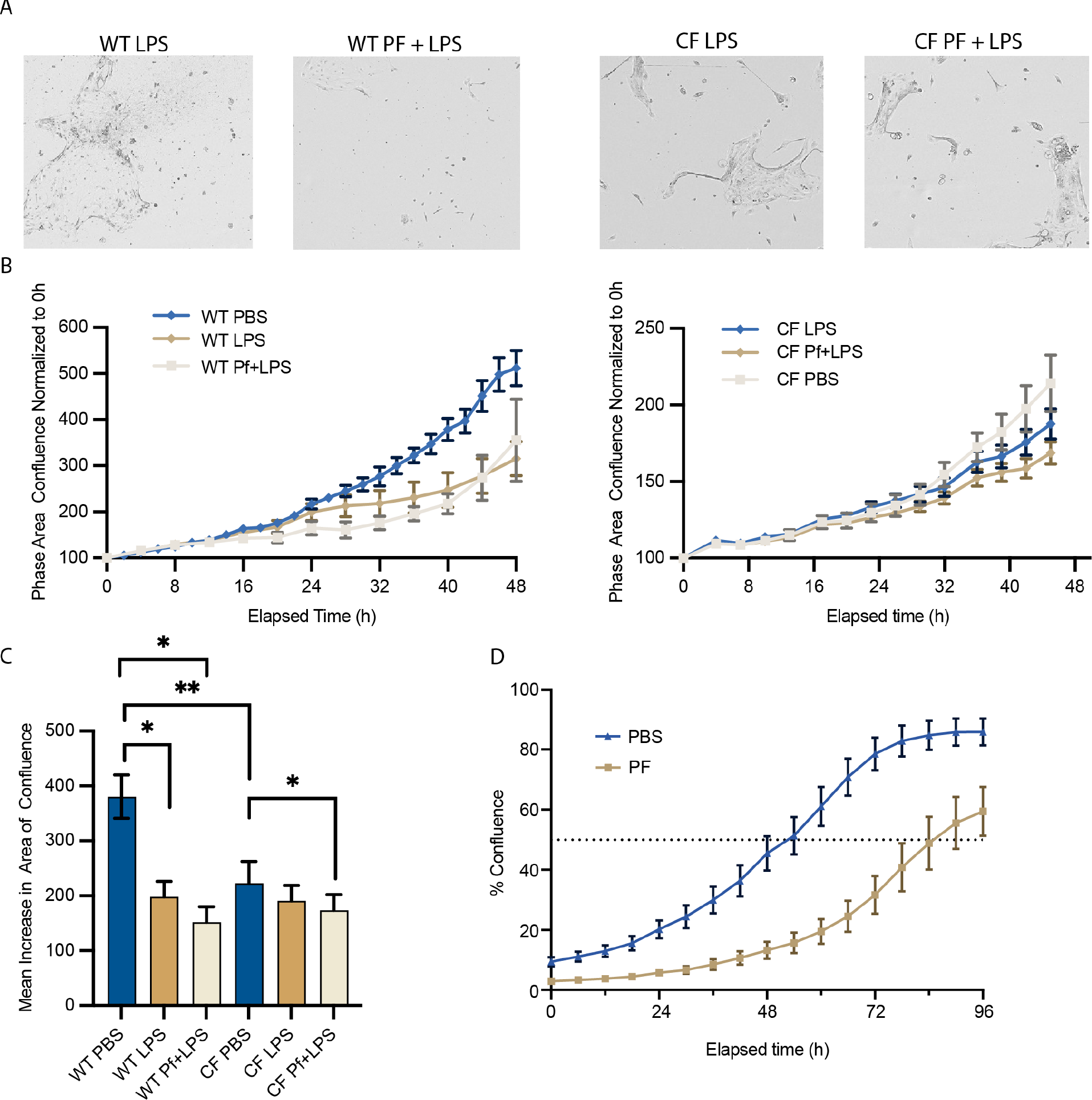
Pf4 phage decreases basal cell progression towards confluence. Basal cell progression towards confluence of the seeded area by proliferation and active migration was tracked for 48hrs via Incucyte. Cells were treated with 1ug/mL LPS ± 10^10^ pfu/mL Pf4 phage at time 0. (A) Representative 200x images of WT cells treated with LPS (left) and Pf+LPS (right) shown at 48hrs. (B) CF cells treated with LPS (left) and Pf+LPS (right) shown at 48hrs. Progression towards confluence shown for WT (C) and CF (D) cells treated with Pf and control conditions. (E) Mean area increase in confluence at 48hrs for both genotypes and all conditions shown. Means and SD estimated by generalized linear model (GLM) and comparisons between different groups assessed by Tukey’s studentized range test. (F) Time to 50% confluence of the seeded area for WT cells in the presence of Pf or PBS control was assessed between 54 and 86 hours (p = 0.012).

These data are in line with the greater magnitude of cluster 3 depletion in CF cells vs. WT cells with Pf treatment. To further delineate the effect on the kinetics of confluence, we extended the period of observation to 96 hours with a second set of similarly treated WT cells. We observed that control WT cells reached 50% confluence by 54 h, as opposed to Pf-treated cells that took 86 h (Fig 6F).

## Discussion

Basal cells are a key member of the airway ecosystem, regenerating multiple cell types in response to injury and infection. We show here that basal cells from both WT individuals and patients with CF exhibit transcriptional heterogeneity in response to LPS. Basal cells segregate into functionally distinct clusters, with the majority of cells falling into four major groups each comprising about 20% of cells in the population. Cluster 0 is defined by GO terms related to neutrophil activation and chemotaxis, indicating a cell population which primarily secretes neutrophil chemoattractants. Cluster 1 is defined by GO terms related to epidermal development and differentiation and likely consists of differentiating basal cells. Cluster 2 is defined by ribosome biogenesis and rRNA processing, indicating this cell grouping is likely undergoing differentiation as well—it is well-established that stem cells maintain low translation rates independently of cell cycle state and dramatically increase ribosome biogenesis during differentiation[24]. Cluster 3 is defined by GO terms relating to positive regulation of cell motility and migration. Two additional minor clusters defined by G1/S cell cycle transitioning (Cluster 4) and cell-substrate junction/hemidesmosome assembly (Cluster 5), comprise a further ∼10% and ∼5% of cells, respectively.

We next investigated the impact of co-stimulating basal cells with LPS to model bacterial infection conditions and a *Pseudomonas*-infecting bacteriophage, Pf4. Our previous work suggests that this bacteriophage is internalized by multiple immune cell types and induces antiviral responses[18], and further work in a cohort of patients with CF indicated that Pf4 phage is associated with worse disease exacerbation and increased chronicity of *Pseudomonas* infection[11]. We find here that basal cells also initiate antiviral immune responses to Pf4 phage. Cluster 6, defined by antiviral genes, increases by 4 and 20-fold, respectively, in WT and CF cell populations. In addition to the increase in cells partitioning to Cluster 6, overall expression of antiviral genes such as IFI6, DDX60, and ISG1is increased overall in both genotypes. Although little work exists on antiviral responses in basal cells, it is known that this cell type can be infected by rhinovirus, and that TLR3 is a major mediator of the airway epithelial cell response to this pathogen[25–27]. Basal cells express TLR3 mRNA, making this a potential mechanism by which Pf phage is sensed in this system. Furthermore, we show that Pf phage, but not LPS alone, specifically induces expression of IRF7, a transcription factor is downstream of TLR3/TRIF signaling primarily induced by type I interferons and viral sensing[28].

We propose that basal cells sense Pf phage through TLR3 and activate an antiviral transcriptional program. A minority of high-expressors drive this antiviral response, but the entire cell population participates to some extent. At the same time, we find that a large subpopulation of basal cells characterized by neutrophil-activating gene expression increases upon Pf phage stimulation. This finding is in line with previous work on viral sensing and neutrophil recruitment: all major neutrophil chemokines are produced during respiratory viral infection, and neutrophil influx to the site of infection is robust and sustained[29]. In particular, TLR3 deficiency has been shown to decrease neutrophil recruitment to the lung, underscoring the involvement of this viral sensor in chemokine production[30]. While Pf phage-exposed basal cells are producing antiviral factors and increasing chemokine production, other cellular functions are neglected. The proportion of cells in Clusters 1-4—corresponding to differentiation, replication, and cell migration—decreased by approximately 5-10% for WT Pf-stimulated cells, and 20-40% for CF cells. We further show that Pf phage exposure decreases the proliferation/migration rate of both WT and CF basal cells.

Taken together, these findings suggest that the presence of this bacteriophage, known to be present in high concentrations (up to 10^11^ copies/mL) in the CF airway, significantly alters the basal cell response to bacterial stimulation. Notably, increased expression of neutrophil activation and chemoattraction factors provides evidence for the basal cell population actively participating in the recruitment of neutrophils to the CF airway. Hyperaccumulation and dysfunction of neutrophils is a known major contributor to airway damage and chronicity of infection in patients with CF [31–33]. In patients colonized with chronic Pf phage-producing *P. aeruginosa*, our results suggest that neutrophil burden would be higher than in the absence of Pf phage, leading to worse patient outcomes in line with our previous work [34]. In parallel, widespread antiviral signaling by basal cells would be expected to promote an aberrant response to Pa infection. Clinically, co-infection with both a viral and bacterial pathogen has been reported to impair antibacterial immune responses: viral infection is known to increase susceptibility to bacterial colonization and decrease control of bacterial replication in mammalian cells [35–38]. Finally, impaired basal cell migration and proliferation in the presence of high Pf phage titer would hinder airway repair and regeneration in the face of chronic bacterial infection.

Although our work identifies a unique and intriguing aspect of *Pa* infection pathobiology in the CF airway, there are limitations to recognize. First, we worked exclusively with nasal biopsies as opposed to bronchial airway cells. While these cells originate from the upper airway, it is well established that *Pa* infection starts in the sinonasal cavity and the nasal epithelium mirrors the pathology and inflammatory response seen in the lower airway[39,40]. Second, our experiments were limited to patients with severe CFTR mutations. Effect on patients with mild mutations or patients receiving CFTR modulators is unclear, and ongoing work is focused on these important aspects.

The findings reported here open several new avenues of investigation. Our focus in this work was on basal epithelial cells, but research into the effects of bacteriophage and LPS stimulation on the fully differentiated airway epithelium would be of great interest. Others have reported that viral infection of basal cells shifts the differentiation trajectory, raising the question of whether chronic Pf phage exposure would have a similar effect [27]. Furthermore, our work was largely limited to a single highly-abundant phage previously known to induce antiviral responses. However, given the resurgence of interest in phage therapy to treat highly antibiotic resistant bacteria in the CF airway and other chronic infection settings, the data presented here suggest that we should not think of bacteriophages as inert, non-immunostimulatory particles. Mammalian cell response to bacteriophages remains an important area of further investigation to identify and reduce unintentional off-target effects in phage therapy.

## Supplemental Figures

**Fig S1:**
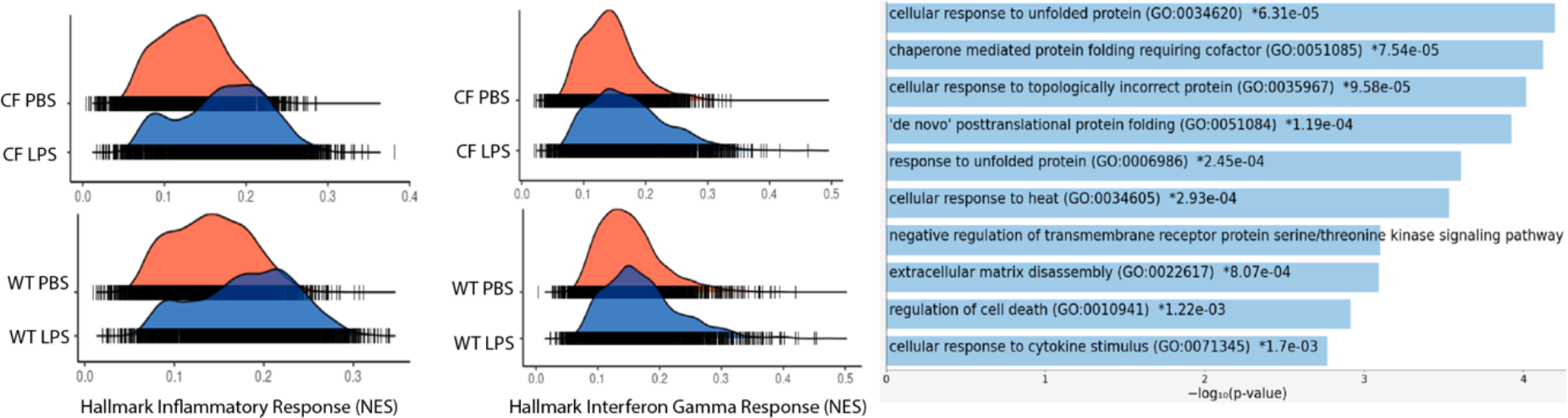
WT and CF Basal Cells exhibit inflammatory LPS responses. (A) GSEA of CF and WT control cell transcriptional response to LPS stimulation. (B) GO enrichment of all significantly differentially expressed genes in CF versus WT control LPS responses.

**Fig S2:**
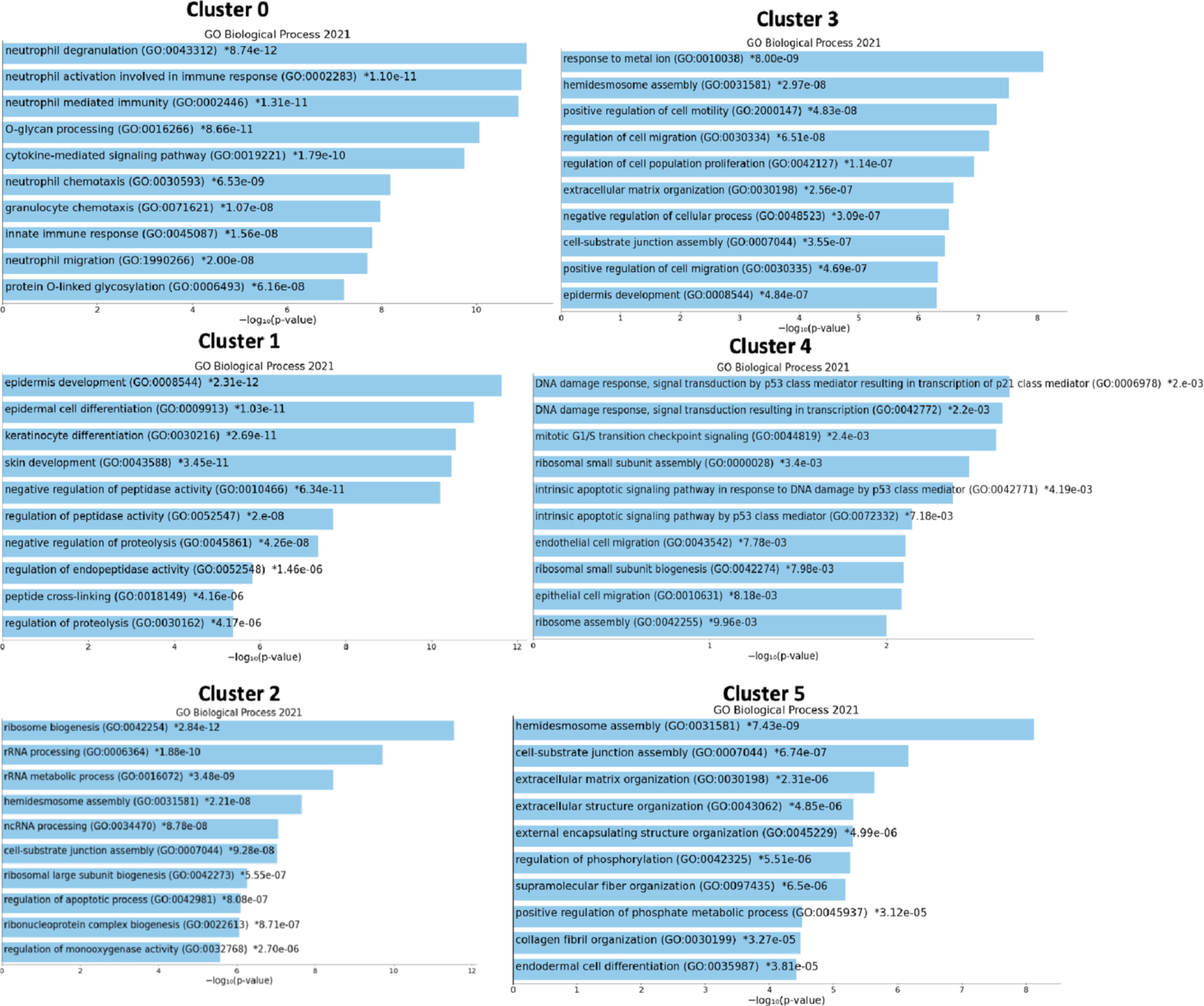
GO analysis of additional clusters from WT and CF cells. GO term enrichment for top upregulated genes in clusters 0-5 for WT and CF donor cells.

**Fig S3:**
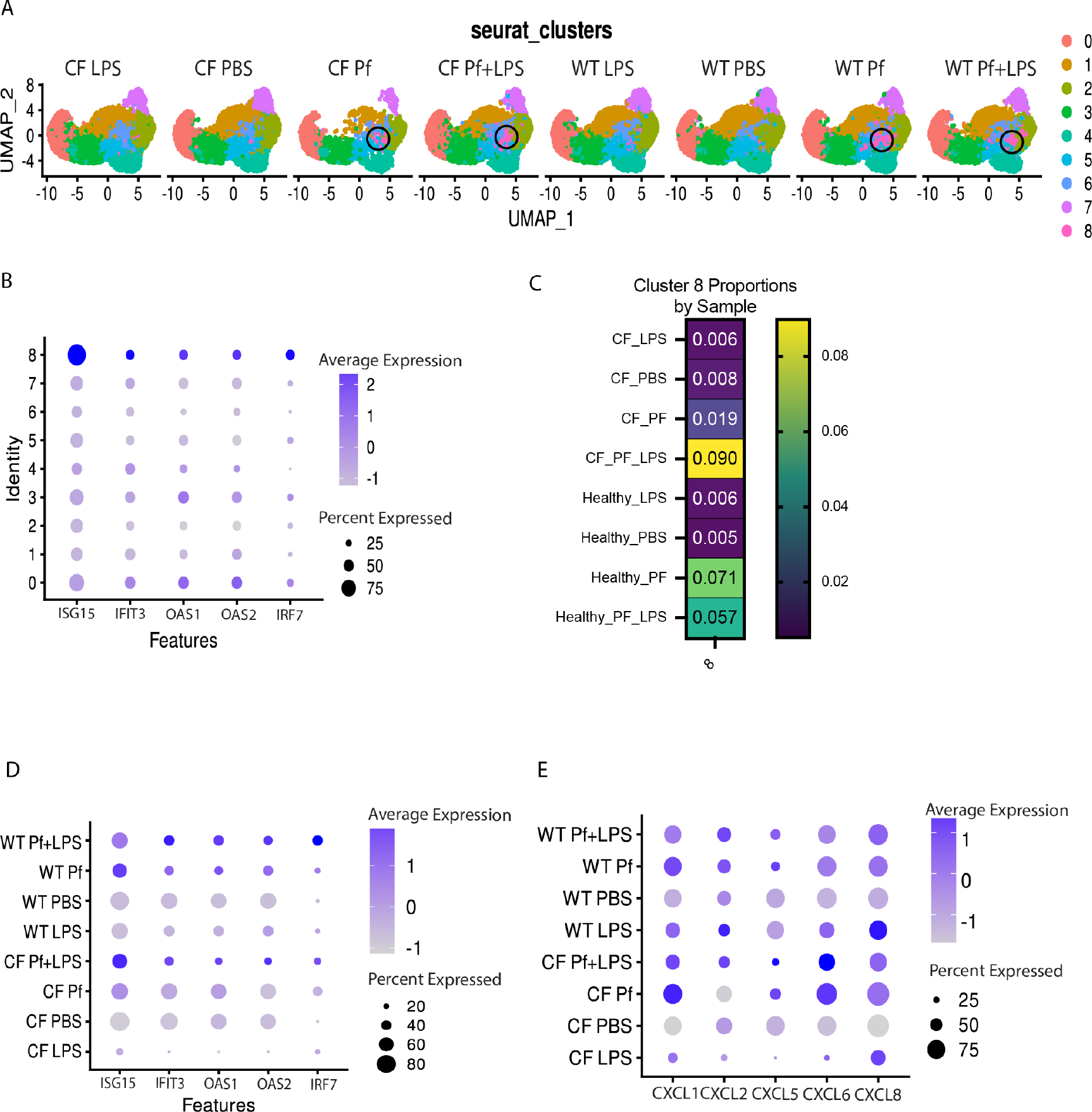
UMAP Clustering with additional controls. A) UMAP plots for all samples, including Pf and PBS only controls for both CF and WT samples. Cluster 8 identi ed corresponding to Cluster 6 of Figure 1B. B) Dot plot showing higher average expression of ISG15, IFIT3, OAS1, OAS2, and IRF7 by cluster. C) Proportion of cells in cluster 8. D) Dot plot showing average expression of markers ISG15, IFIT3, OAS1, OAS2, and IRF7 across samples including Pf and PBS controls. E) Dot plot showing average expression of markers CXCL1, CXCL2, CXCL5, CXCL6, and CXCL8 across all samples including Pf and PBS controls

**Fig S4:**
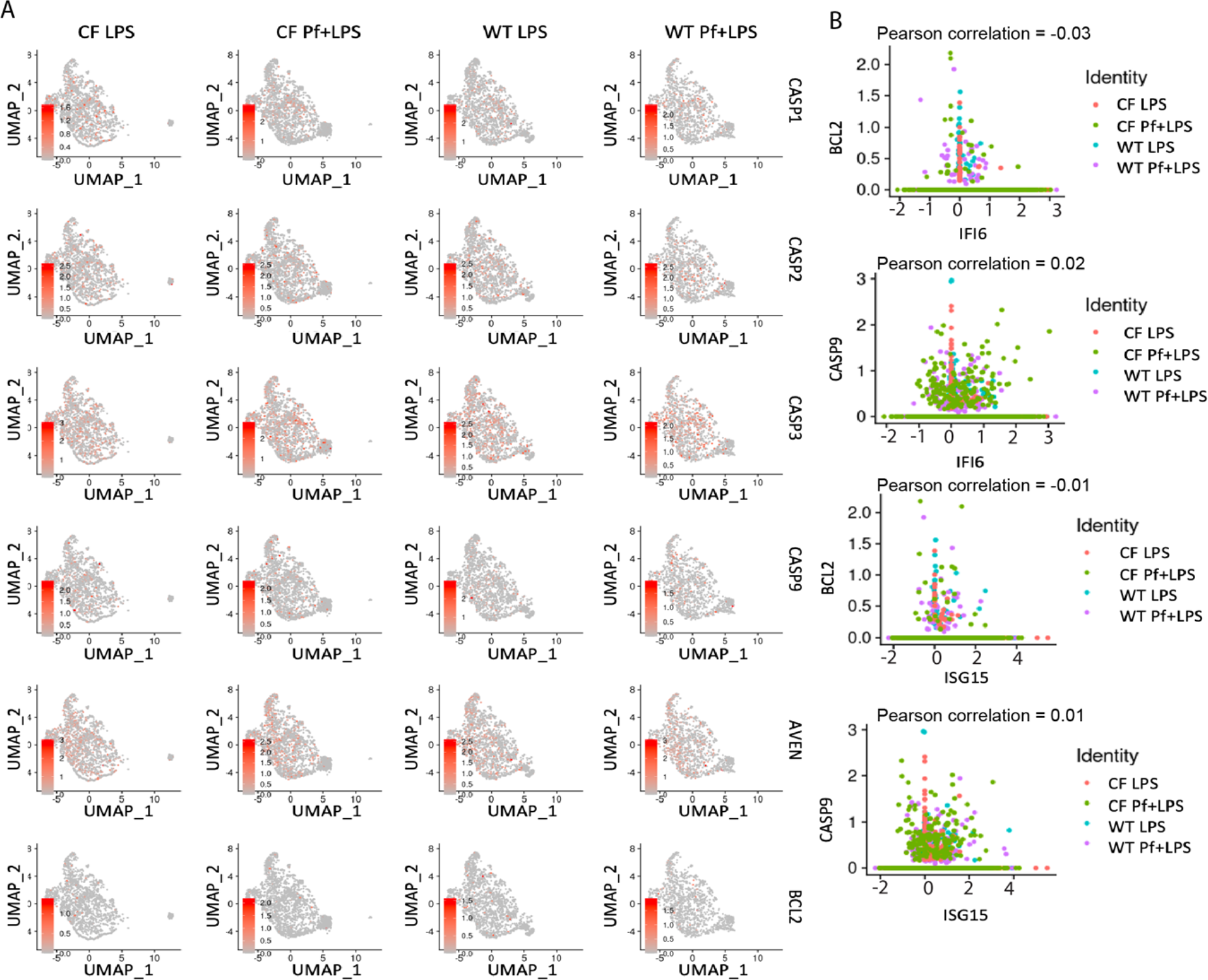
Feature plots of apoptotic markers in basal cells. (A) Ridge plots for apoptosis markers CASP1, CASP2, CASP3, CASP9, AVEN, and BCL2. (B) Feature Scatter plots for BCL2 and CASP9 versus antiviral signature genes ISG15 and IFI6, with pearson correlation values above plot.

**Fig S5:**
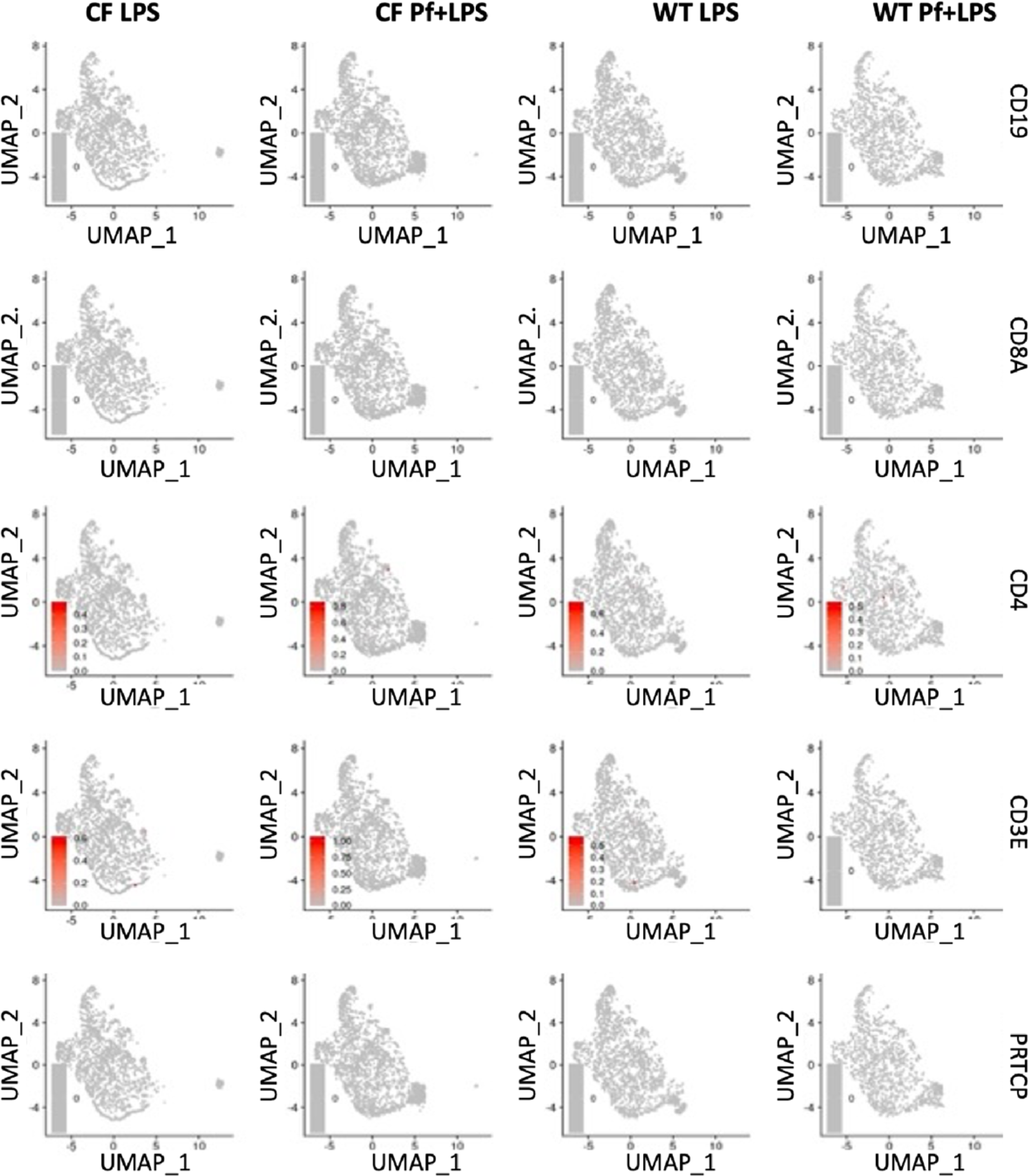
Feature plots of lymphocyte markers in WT and CF cells. Ridge plots for immune cell markers CD19, CD8A, CD4, CD3E, and PTPRC (CD45) in WT and CF donor cells.

